# Discovery of TDP-43 aggregation inhibitors *via* a hybrid machine learning framework

**DOI:** 10.64898/2026.02.12.705375

**Authors:** Sofia Kapsiani, Sulay Vora, Ana Fernandez-Villegas, Clemens F. Kaminski, Nino F. Läubli, Gabriele S. Kaminski Schierle

## Abstract

TAR DNA-binding protein 43 (TDP-43) aggregation is a hallmark of several neurodegenerative diseases, including amyotrophic lateral sclerosis and frontotemporal dementia. Recent therapeutic efforts have highlighted the potential of small molecules capable of inhibiting TDP-43 aggregation; however, no effective treatments currently exist. Here, we developed a hybrid machine learning approach combining graph neural network (GNN) embeddings with traditional chemical descriptors and biological target annotations. Using XGBoost as the final classifier enabled model interpretability through SHAP analysis, allowing the identification of key chemical features and target annotations associated with TDP-43 anti-aggregation activity. Complementary Monte Carlo Tree Search analysis highlighted specific chemical substructures linked to predicted activity. By screening an external library of 3,853 small molecules, the model identified two compounds not previously evaluated against TDP-43 aggregation, namely berberrubine and PE859. Molecular docking analysis revealed that both compounds interact favourably with the TDP-43 RNA recognition motif (RRM) domain through distinct binding modes. Experimental validation showed that both compounds significantly reduced TDP-43 aggregation in HEK cells. Further testing in *Caenorhabditis elegans* expressing human TDP-43 demonstrated that PE859 significantly rescued locomotor defects, while berberrubine showed partial improvement. This work establishes a hybrid machine learning approach for accelerating small molecule drug discovery, yielding two promising therapeutic candidates for TDP-43 proteinopathies.

## 1 Introduction

Neurodegenerative diseases are characterised by the progressive loss of neuronal function, ultimately leading to cell death[1, 2]. While clinical symptoms vary across these disorders, they share a common pathological hallmark, the abnormal aggregation of disease-associated proteins[1, 3]. One such protein is TAR DNA-binding protein 43 (TDP-43), which is implicated in several of these diseases, including amyotrophic lateral sclerosis (ALS), frontotemporal dementia (FTD), and Alzheimer’s disease[4]. Specifically, in healthy cells, TDP-43 is primarily located in the nucleus, where it contributes to gene expression and RNA processing and regulation[4, 5]. However, under disease-related conditions, TDP-43 undergoes post-translational modifications, such as cleavage, ubiquitination, and phosphorylation, that lead to its mislocalisation from the nucleus to the cytoplasm and the formation of insoluble aggregates[4, 5]. This mislocalisation results in the loss of TDP-43’s normal physiological function, with both the loss of function and the gain of toxic function believed to contribute to disease pathogenesis[6].

Although a limited number of small molecule candidates targeting TDP-43 pathology have progressed to clinical trials, there is currently no effective therapy for slowing or reversing TDP-43 proteinopathies[7]. In an effort to accelerate therapeutic discovery, machine learning has become an increasingly valuable tool for identifying drug candidates across a range of biomedical applications, including Alzheimer’s disease[8–10], Parkinson’s disease[11–13], and ALS[14, 15]. Notably, Gao *et al*. (2023)[16] applied deep learning using graph convolutional networks to identify small molecules that reduce pathological TDP-43 nuclear liquid–liquid phase separation (LLPS), demonstrating the efficiency of machine learning-based drug screening[16]. Here, we focus on predicting inhibitors of TDP-43 aggregation rather than LLPS and go one step further by integrating chemical descriptors and biological target annotations with molecular graph embeddings to develop a machine learning framework that enables model interpretability.

After training and evaluation, we applied our model to screen an external dataset of small molecules to identify novel TDP-43 aggregation inhibitors. From the compounds predicted with high probability of being active (*>*80 %), two high-confidence candidates, namely berberrubine and PE859, were selected for experimental evaluation. Both compounds significantly reduced TDP-43 aggregation in human embryonic kidney (HEK) cells, as measured by fluorescence lifetime imaging microscopy. Furthermore, in *Caenorhabditis elegans (C. elegans)* expressing wild-type human TDP-43 pan-neuronally, PE859 significantly ameliorated locomotor deficits, whereas berberrubine exhibited a partial improvement.

## 2 Results and discussion

### 2.1 A hybrid machine learning framework for predicting TDP-43 aggregation inhibitors

To predict TDP-43 anti-aggregation activity, we employed a hybrid modelling approach integrating a graph neural network (GNN) with an XGBoost (eXtreme Gradient Boosting)[17]. This approach leverages the GNN’s ability to learn task-specific molecular representations[18] with the interpretability and robustness of decision tree-based algorithms[19]. First, a directed message-passing neural network (D-MPNN), implemented using Chemprop[20], was trained to generate molecular embeddings that integrate localised atomic and bond-level information. These molecular embeddings were then extracted and combined with two additional feature types (Figure 1a), (i) chemical descriptors computed using the RDKit Python library[21] to encode global physicochemical and topological properties, and (ii) biological target information obtained from ChEMBL[22] to provide insights into known protein interactions. The combined features were used to train an XGBoost classifier, as shown by Figure 1b, which was selected as the final classifier due to the relatively small size of the dataset and the substantial class imbalance (10.6% active), conditions under which decision tree–based approaches have demonstrated strong performance while providing high interpretability[19, 23].

**Figure 1.**
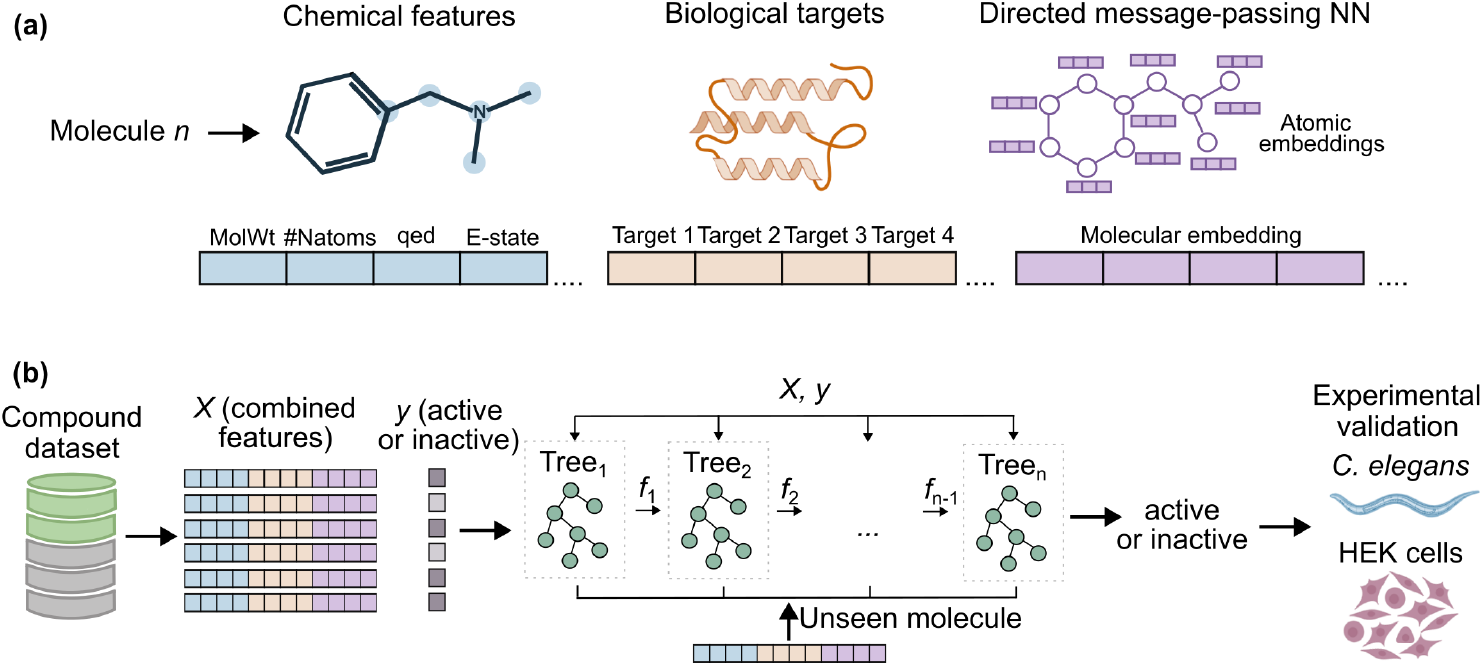
Hybrid modelling approach for predicting TDP-43 aggregation inhibitors. (a) The model uses a combination of feature types, including chemical descriptors (e.g., molecular weight, number of nitrogen atoms), ChEMBL biological target annotations, and GNN-derived molecular embeddings as an input. (b) The combined feature set is used to train an XGBoost classifier to predict whether a compound is active as a TDP-43 aggregation inhibitor. The model is applied to an external screening dataset, and high-confidence hits were experimentally validated in *C. elegans* and Human Embryonic Kidney (HEK) cells.

The model was trained on a manually curated dataset comprising 294 small molecules previously reported to protect against or reduce TDP-43 aggregation and 2,482 molecules lacking such activity. Data were collected from literature, ChEMBL[22], PubChem[24], and patent records, encompassed molecules tested across diverse biological systems such as cell lines and model organisms.

The dataset was split into an 80% training set and a 20% test set. On the training set,5-fold nested cross-validation was performed for hyperparameter optimisation and feature selection. Cross-validation folds were defined using Butina clustering[25], such that structurally similar compounds were assigned to the same fold, enabling evaluation on structurally distinct molecules and assessing generalisation to novel chemical scaffolds[26]. Feature selection involved first removing redundant and highly correlated features to reduce noise, followed by Recursive Feature Elimination (RFE)[27] to select the most informative features. Following cross-validation, a final model was trained on the full training set using the optimal hyperparameters identified during cross-validation, and feature selection was repeated using the same protocol. Model performance was then evaluated on the test set.

### 2.2 Combined GNN embeddings and traditional molecular features improve performance for predicting TDP-43 aggregation inhibitors

To determine which features best predict TDP-43 aggregation inhibitors, models were trained using individual feature types and their combinations. Model performance was evaluated using metrics suitable for imbalanced datasets[28]: Area Under the Receiver Operating Characteristic Curve (ROC-AUC), F1-score, Precision, Matthews Correlation Coefficient (MCC), and Balanced Accuracy.

Among single-feature models, biological targets from ChEMBL outperformed the sole use of chemical descriptors, extended-connectivity fingerprints (ECFPs, 2,048-bit length), or conventional GNN with a Multilayer Perceptron (MLP) classifier in both cross-validation (Supplementary Table 1) and test set evaluation (Table 1).

**Table 1.** Performance metrics on the test set for models trained using individual and combined feature types.

| Features | Classifier | ROC-AUC | MCC | F1-score | Balanced Accuracy | Precision |
| --- | --- | --- | --- | --- | --- | --- |
| GNN embeddings + chemical descriptors + biological targets | XGBoost | 0.84 | **0.42** | **0.37** | **0.61** | **0.87** |
| ECFPs (2,048) chemical descriptors + biological targets | XGBoost | **0.88** | 0.29 | 0.22 | 0.56 | 0.78 |
| chemical descriptors + biological targets | XGBoost | **0.88** | 0.39 | 0.34 | 0.60 | 0.80 |
| biological targets | XGBoost | 0.87 | 0.34 | 0.27 | 0.58 | 0.82 |
| chemical descriptors | XGBoost | 0.81 | 0.06 | 0.03 | 0.51 | 0.33 |
| GNN | MLP | 0.78 | 0.16 | 0.10 | 0.52 | 0.60 |
| ECFPs (2,048) | XGBoost | 0.72 | 0.04 | 0.08 | 0.51 | 0.17 |
| random Y shuffle (GNN embeddings + chemical descriptors + biological targets) | XGBoost | 0.53 | 0.00 | 0.00 | 0.50 | 0.00 |
| GNN embeddings + chemical descriptors + biological targets | Random Forest | 0.81 | 0.20 | 0.13 | 0.53 | 0.67 |

Importantly, the combined model integrating GNN embeddings, chemical descriptors, and biological targets achieved the strongest overall performance, with the highest F1-score, MCC, and Balanced Accuracy in cross-validation. On the test set, the combined model achieved the highest F1-score, MCC, Balanced Accuracy, and Precision (Table 1). While ROC-AUC was slightly lower than some other models, this metric evaluates performance across all classification thresholds (Figure 2a) rather than at a fixed decision boundary, and is sensitive to class imbalance[29].

**Figure 2.**
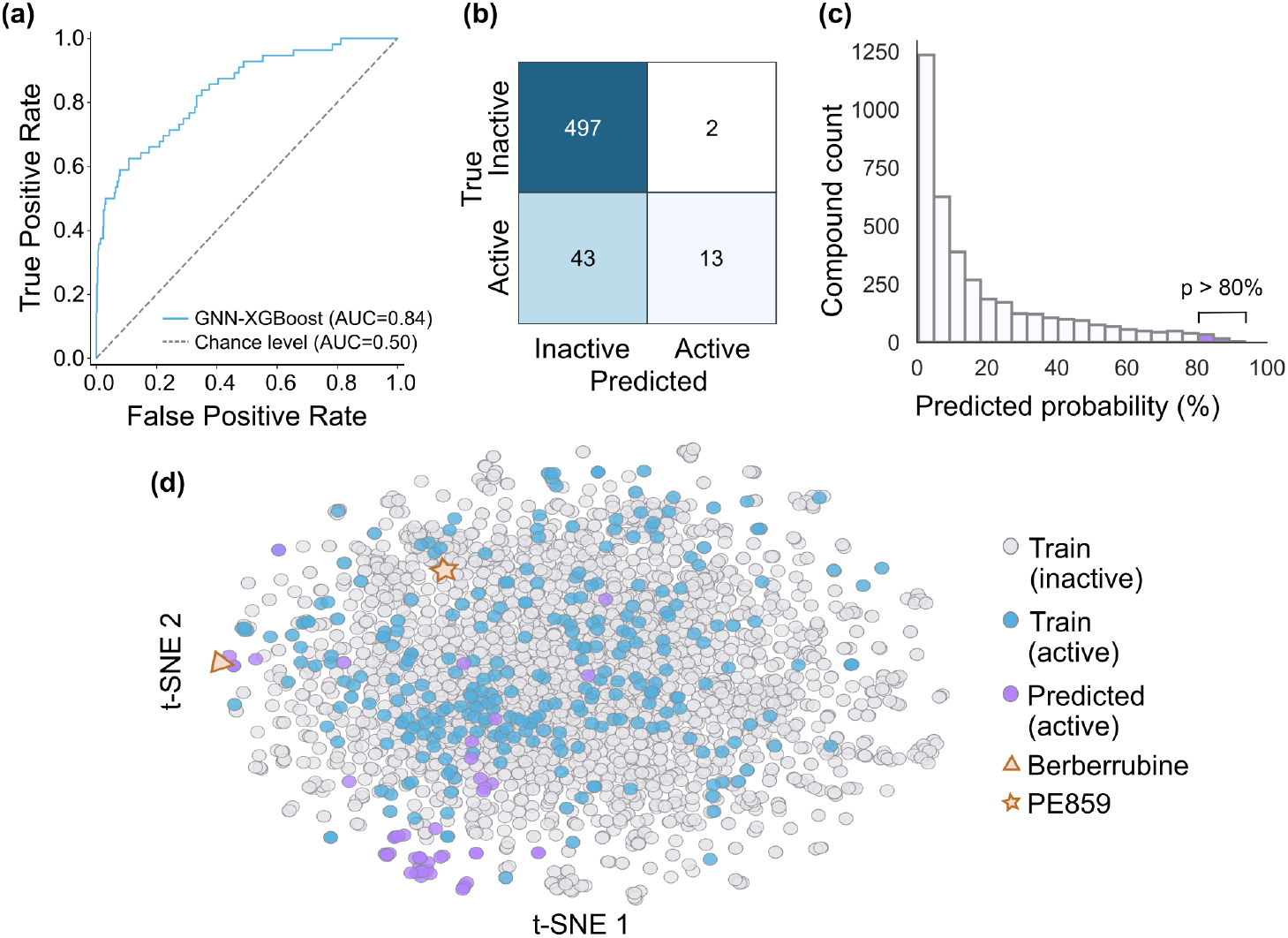
Model performance on held-out test set and computation screening of external dataset. (a) ROC curve of the GNN-XGBoost model on the held-out test set, shown with a blue line (AUC = 0.84), and the chance level shown as a dashed grey line (AUC = 0.50). (b) Confusion matrix showing model performance on the test set. (c) Histogram of predicted probabilities for the external screening dataset; subsequent analysis focused on compounds with a predicted probability above 80%. (d) Chemical space visualisation using t-distributed stochastic neighbour embedding (t-SNE) applied to 2,048-bit extended-connectivity fingerprints (ECFPs). Each dot represents a single compound, with grey indicating inactive training compounds, blue indicating active training compounds, and purple indicating external compounds with a predicted probability above 80%. Berberrubine and PE859, selected for experimental validation, are highlighted with a triangle and a star in orange, respectively

As shown in the confusion matrix (Figure 2b), the combined model (GNN embeddings + chemical descriptors + biological targets) predicted 15 compounds as active on the test set, of which 13 were true positives. This indicates that compounds predicted as active have a high probability of having TDP-43 anti-aggregation activity. However, the model misclassified 43 true actives as inactive, reflecting a higher false-negative rate. Here, this trade-off is acceptable, as minimising false positives is critical to reducing experimental cost and time.

To ensure that model’s performance was not due to chance or an information leakage, we further trained a control model with randomly shuffled labels in the training set. As expected, this control model performed at chance level (Table 1), validating the integrity of the modelling pipeline. Finally, we demonstrated that using XGBoost as the final classifier yielded improved performance compared to Random Forest[30], another widely used tree-based classification approach (Table 1).

### 2.3 Virtual screening of an external dataset identifies novel TDP-43 aggregation inhibitors

The combined model was applied to screen an external library of 3,853 small molecules, compiled from the work of Smer-Barreto *et al*.(2023)[31]. The distribution of predicted probabilities of the external dataset compounds is shown in Figure 2c. Our analysis focused on compounds with a predicted probability *>* 80% of being active, with a total of 57 molecules meeting this threshold. The chemical space of these top candidates, alongside the training set compounds, was visualised using the dimensionality reduction technique t-distributed stochastic neighbour embedding (t-SNE) applied to 2,048-bit ECFPs (Figure 2d). The top 57 hits were then further filtered using criteria prioritising oral bioavailability (Lipinski’s Rule of Five), blood–brain barrier permeability, limited structural alerts, and no prior testing against TDP-43 aggregation.

Finally from these filtered candidates, two compounds were selected for experimental evaluation. Berberrubine, the top hit from the external dataset with a predicted probability of 94%, is a metabolite of berberine, a known active compound from the training set[32]. It was prioritised to validate the chemical-space cluster containing other highly ranked predicted compounds, including Sanguinarine, Nitidine chloride, Coptisine chloride, Cepharanthine, and Epiberberine. Additionally, PE859 (predicted probability of 83%) was selected due to its lower structural similarity (Tanimoto coefficient 0.33) to known actives, representing a more novel chemical scaffold within the chemical space.

### 2.4 Explainable machine learning analysis identifies key chemical and biological features

To identify features most strongly influencing TDP-43 anti-aggregation predictions, a SHAP (SHapley Additive exPlanations)[33] analysis was performed. Of the 30 top-ranked features, (shown in Supplementary Figure 1) half were GNN-derived embeddings, highlighting the relevance of the selected hybrid approach. However, as GNN-derived embeddings themselves are abstract and not directly interpretable, further analysis was focused on the non-GNN features (Figure 3a).

**Figure 3.**
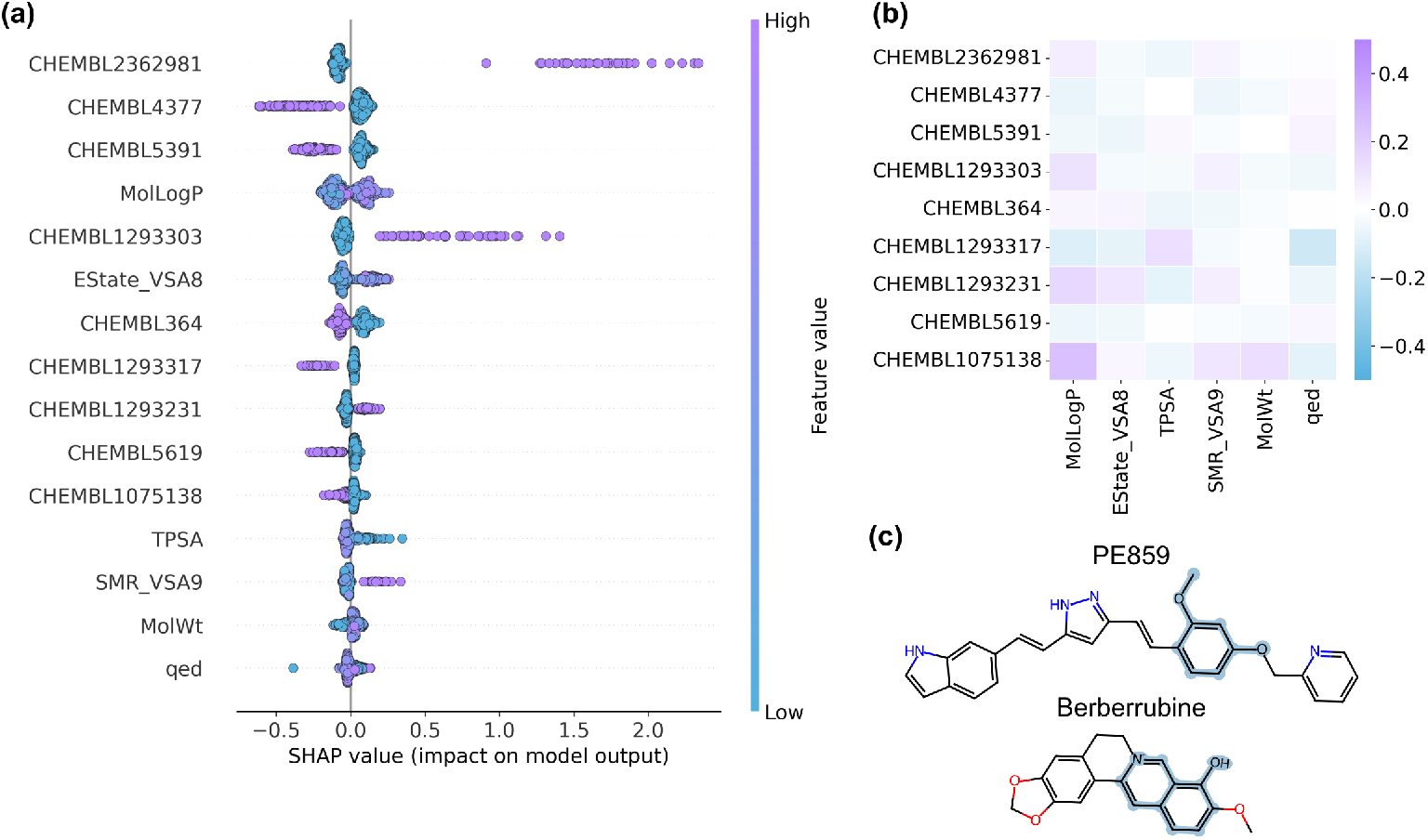
Identification of features that influence model predictions. (a) SHAP beeswarm plot showing the 15 highest-ranked non-GNN features (y-axis). GNN-derived embeddings were excluded due to their limited interpretability. Each point represents the SHAP value of a single test set compound for that feature, coloured by its corresponding feature value (blue, low; purple, high)[34]. SHAP values on the x-axis indicate how high or low feature values shift predictions towards the active or inactive class[35]. (b) Correlation matrix between top-ranked biological targets and chemical descriptors, showing only weak associations, and suggesting that these features capture complementary information. (c) Chemical structures of compounds selected for experimental validation (PE859 and berberrubine), with rationales identified by Monte Carlo Tree Search highlighted in blue.

Among chemical descriptors, increased lipophilicity (MolLogP)[36] and reduced polar surface area (TPSA)[37] correlated with higher predicted activity, consistent with blood–brain barrier penetration requirements[38]. Similarly, higher molecular surface area descriptors (ES-tate VSA8, SMR VSA9), reflecting electronic[39] and polarisability[36] properties, also associated with increased activity, with polarisability having previously been shown to facilitate interactions with amyloidogenic proteins[40]. Finally, increased molecular weight (MW) and reduced drug-likeness (QED)[41] favoured the active class, potentially reflecting a dataset bias arising from the presence of structurally complex natural products among the active training compounds.

Regarding biological targets, several annotations showed positive associations with predicted activity. The interaction with CHEMBL2362981 (TDP-43) had the strongest positive association, although this annotation is not solely determinative of activity, as it appears for both active (∼ 30%) and inactive (∼ 3.4%) compounds. Interaction with CHEMBL1293231 (ROR-*γ*), a regulator of inflammatory signalling[42], also displayed a positive association. In contrast, predicted interactions with CHEMBL4377 (G*α*s), CHEMBL5391 (DNA polymerase *ι*), CHEMBL5619 (APEX1), and CHEMBL1075138 (TDP1) were associated with reduced predicted activity, potentially reflecting the model’s tendency to penalise compounds interacting with DNA-repair or G-protein signalling pathways. The model also incorporated information from other RNA-binding proteins, e.g., CHEMBL1293303 (Nonstructural protein 1, positive association) and CHEMBL1293317 (Muscleblind-like protein 1, negative association). While the applied SHAP analysis highlighted target annotations that influenced predictions, it remains unclear whether these associations reflect biological mechanisms or chemical features shared by compounds annotated with these targets. A subsequent correlation analysis (Figure 3b) further revealed only weak associations between top-ranked target annotations and chemical descriptors, thus suggesting that these features are able to capture complementary information.

Although GNN embeddings cannot be directly mapped to chemical substructures, Monte Carlo Tree Search analysis[43] identified graph-based “rationales” that shift predictions of the pretrained GNN towards the active class. While the final predictive model used the precomputed GNN embeddings rather than the GNN itself, this approach remains informative for identifying structural motifs associated with predicted activity, with the analysis highlighting aromatic groups enriched with heteroatoms (O, N, S) as key motifs (see Supplementary Table 2). Rationales for compounds prioritised for experimental testing, berberrubine and PE859, are shown in Figure 3c.

**Table 2.**
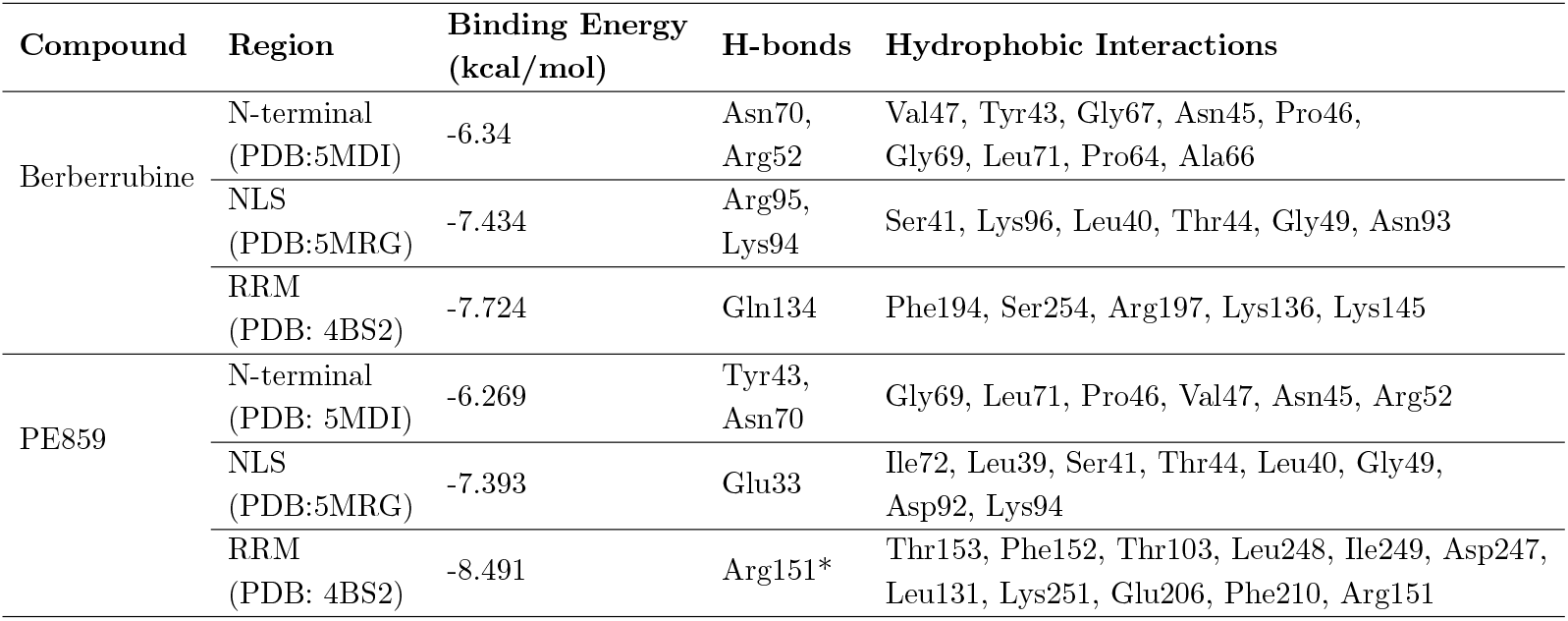
Docking simulations were performed against the N-terminal domain (PDB IDs: 5MDI, 5MRG) and the RNA recognition motif (RRM; PDB ID: 4BS2) of TDP-43. Binding energies (kcal/mol) correspond to the top-ranked poses. Ligand–residue interactions were identified using LigPlus [44], with asterisked hydrogen bonds (*) confirmed by manual inspection in PyMOL[45].

| Compound | Region | Binding Energy (kcal/mol) | H-bonds | Hydrophobic Interactions |
| --- | --- | --- | --- | --- |
| Berberrubine | N-terminal (PDB:5MDI) | -6.34 | Asn70, Arg52 | Val47, Tyr43, Gly67, Asn45, Pro46, Gly69, Leu71, Pro64, Ala66 |
|  | NLS (PDB:5MRG) | -7.434 | Arg95, Lys94 | Ser41, Lys96, Leu40, Thr44, Gly49, Asn93 |
|  | RRM (PDB: 4BS2) | -7.724 | Gln134 | Phe194, Ser254, Arg197, Lys136, Lys145 |
| PE859 | N-terminal (PDB: 5MDI) | -6.269 | Tyr43, Asn70 | Gly69, Leu71, Pro46, Val47, Asn45, Arg52 |
|  | NLS (PDB:5MRG) | -7.393 | Glu33 | Ile72, Leu39, Ser41, Thr44, Leu40, Gly49, Asp92, Lys94 |
|  | RRM (PDB: 4BS2) | -8.491 | Arg151* | Thr153, Phe152, Thr103, Leu248, Ile249, Asp247, Leu131, Lys251, Glu206, Phe210, Arg151 |

### 2.5 Molecular docking predicts favourable interactions of Berberrubine and PE859 with TDP-43 RNA-recognition domains

Next, we performed molecular docking to predict the potential interactions of the selected compounds, berberrubine and PE859, with TDP-43 across three functionally relevant regions. Specifically, we investigated binding energies at the N-terminal, the nuclear localisation sequence (NLS), and the RNA recognition motif (RRM) region (RRM1–RRM2), with both compounds demonstrating favourable binding energies across all regions as shown in Table 2.

The highest predicted affinities were observed within the RRM region, with the compounds occupying distinct binding pockets as shown in Figures 4a and 4b. Specifically, berberrubine docked at the RRM1–RRM2 interface (*−*7.72 kcal/mol), with predicted interactions involving Gln134, Phe194, Ser254, Arg197, Lys136, and Lys145 (Figure 4c). This site partially overlaps with a previously reported adenosine triphosphate (ATP) binding pocket[46], which has been implicated in the inhibition of RRM aggregation[47]. Similarly, PE859 exhibited the strongest overall binding affinity in the RRM region (*−*8.49 kcal/mol), where it formed a predicted hydrogen bond with Arg151 (Figure 4d), as well as extensive hydrophobic interactions with multiple residues, including Asp247 (Table 2). This binding site is of particular interest, as Arg151 and Asp247 form a critical salt bridge that contributes to RNA recognition and the structural stability of TDP-43[46].

**Figure 4.**
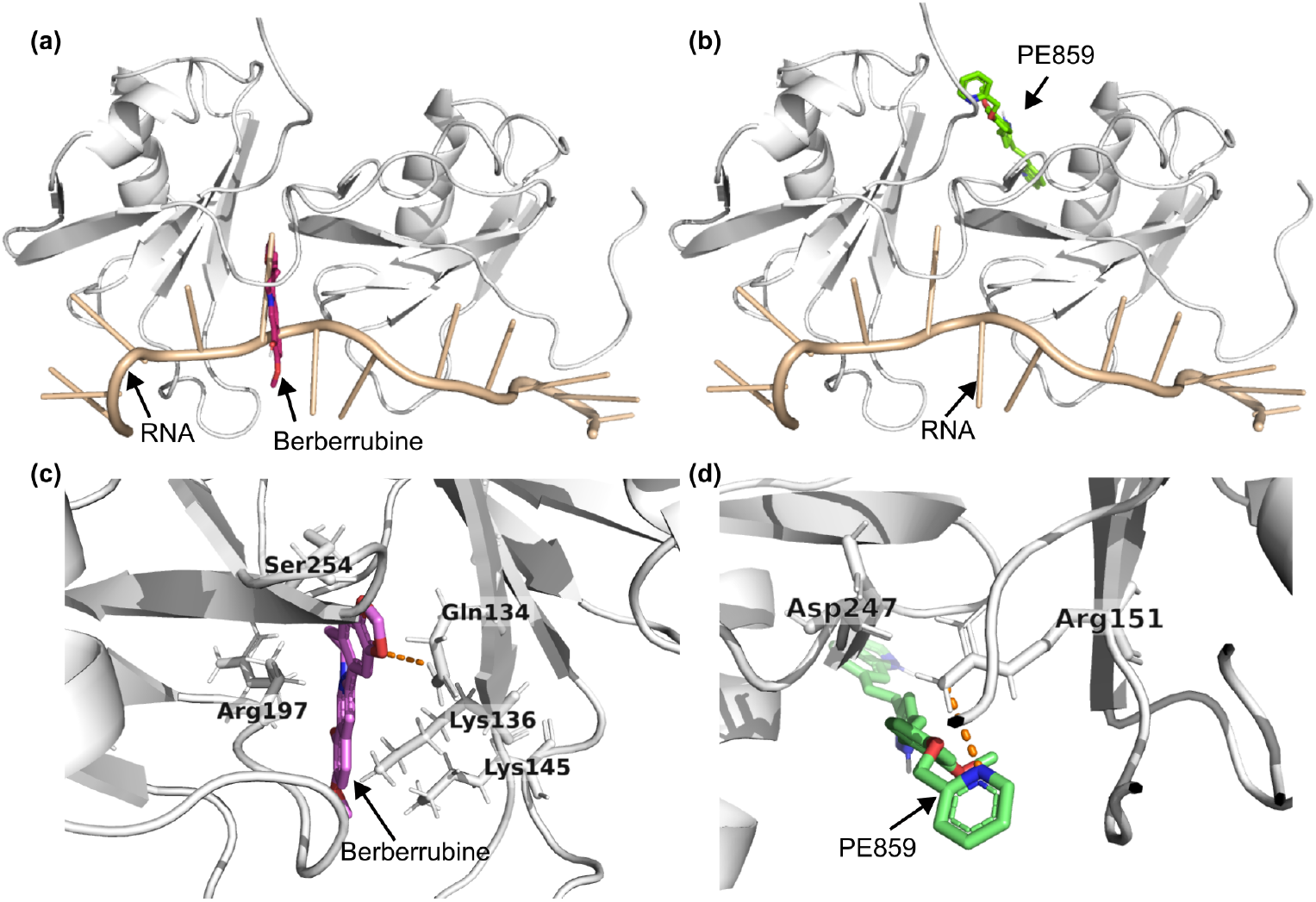
Molecular docking predicts distinct binding modes of berberrubine and PE859 within the TDP-43 RRM domain. Docking poses of (a) berberrubine and (b) PE859 in the RNA recognition motif (RRM; PDB ID: 4BS2) of TDP-43. The RRM domain is shown in light grey, and bound RNA in light orange. (c) Berberrubine was predicted to bind at the interface between RRM1 and RRM2, whereas (d) PE859 binds in close proximity to the Arg151–Asp247 salt bridge, forming a hydrogen bond with Arg151, as confirmed by manual inspection in PyMOL. Hydrogen bonds are shown as orange dashed lines.

### 2.6 Berberrubine and PE859 reduce TDP-43 aggregation in HEK cells and improve *C. elegans* locomotion

Berberrubine and PE859 were further evaluated in HEK cells. Cell viability was first assessed using an MTS assay to determine suitable working concentrations. PE859 exhibited greater cytotoxicity than berberrubine, leading to the selection of 10 *µ*M berberrubine and 5 *µ*M PE859 to balance efficacy with cell viability (Supplementary Figures 2a and 2b).

Then, fluorescence lifetime imaging microscopy (FLIM) was used to examine the effects of the compounds in HEK cells expressing GFP-tagged TDP-43. Changes in GFP fluorescence lifetime served as a readout of TDP-43 aggregation, as aggregation reduces the distance between the GFP molecules, thereby increasing the fluorescence self-quenching and inducing a lower fluorescence lifetime[48]. Importantly, the measurements revealed that treatment with either berberrubine or PE859 significantly increased GFP lifetime compared with the DMSO vehicle control (Figures 5a and b), thus indicating that the compounds were able to reduce *in cellulo* aggregation of TDP-43.

**Figure 5.**
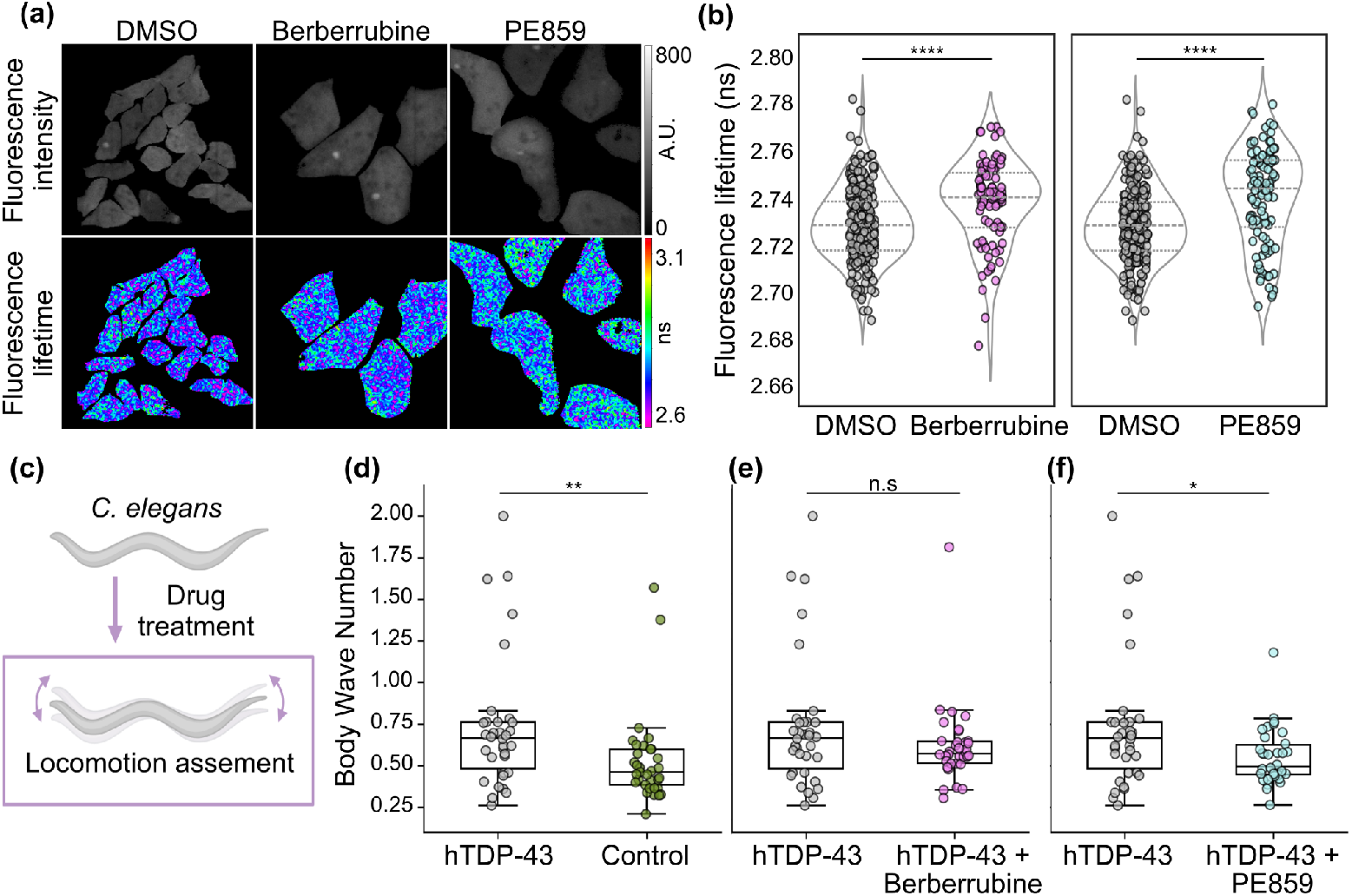
Berberrubine and PE859 reduce GFP–TDP-43 aggregation and improve locomotor defects. (a) Fluorescence intensity (top) and modulation fluorescence lifetime (bottom) images of GFP–TDP-43 expressed in HEK cells treated with DMSO vehicle control (left), 10 *µ*M berberrubine (middle), or 5 *µ*M PE859 (right). (b) Violin plots showing modulation fluorescence lifetime values of GFP–TDP-43 in HEK cells treated with DMSO vehicle control compared with 10 *µ*M berberrubine (left) or 5 *µ*M PE859 (right). Data represent *≥* 3 biological replicates per condition; each dot corresponds to an individual cell (n *≥* 88 cells per condition). Statistical analysis was performed using a two-tailed Student’s t-test for berberrubine and Welch’s t-test for PE859. (c) *C. elegans* assay used to assess rescue of TDP-43–induced locomotor defects following drug treatment. Boxplots show body wave number for worms expressing human TDP-43 compared with (d) control worms, (e) TDP-43 worms treated with 10 *µ*M berberrubine, and (f) TDP-43 worms treated with 10 *µ*M PE859. Data were collected over *≥* 3 biological repeats, with at least 33 worms analysed per condition. Statistical analysis was performed using the Mann–Whitney U test, with *p *<* 0.05, **p *<* 0.01, ****p *<* 0.0001; n.s., not significant.

Next, the compounds were evaluated *in vivo* by assessing their ability to rescue TDP-43 induced locomotor defects in *C. elegans* nematodes. Specifically, two strains i.e. control worms and worms pan-neuronally expressing human TDP-43 (hTDP-43), were analysed, and their locomotion was quantified using a swimming assay (Figure 5c) in combination with computational feature extraction. Worms expressing hTDP-43 exhibited a significantly increased body wave number compared with control worms (Figure 5d), consistent with previous reports that elevated body wave number has been associated with ageing and disease-related locomotion impairment[49]. However, following treatment with 10 *µ*M berberrubine produced a partial, non-significant reduction in body wave number (Figure 5e), whereas 10 *µ*M PE859 significantly decreased body wave number relative to untreated hTDP-43 worms (Figure 5f), indicating a stronger functional rescue.

Analysis of travel speed, another locomotion parameter previously shown to decrease in *C. elegans* models of neurodegeneration[50], revealed that, while TDP-43 expression produced a modest, non-significant reduction compared to the control condition, both berberrubine and PE859 significantly increased travel speed relative to untreated TDP-43 worms (Supplementary Figure 3). Together, these data suggest that both compounds improve locomotor function in TDP-43-expressing worms, with PE859 demonstrating more robust rescue across multiple locomotor parameters.

## 3 Conclusion

We present a hybrid machine learning framework that integrates GNN-derived embeddings with classical chemical descriptors and biological target annotations to predict small molecule inhibitors of TDP-43 aggregation. Explainability analyses indicated that predicted activity is associated with features linked to blood–brain barrier penetration, including higher lipophilicity and lower polar surface area[38], as well as larger, more complex natural-product-like scaffolds, which may reflect biases in publicly available data. The model disfavors compounds targeting DNA repair pathways, which may help minimise off-target toxicity, and shows a positive association with ROR-*γ*, consistent with the established role of inflammation in neurodegeneration[3]. While, GNN-derived embeddings were the most prominent features for classification, a limitation is that they are inherently difficult to interpret.

Application to an external compound dataset identified multiple, high-confidence candidates with reported neuroprotective activity. Top-ranked compounds, including sanguinarine, Tubeimoside I, and ketoconazole, have previously been reported to reduce TDP-43 aggregation[51–53], thereby providing retrospective validation. Future work could evaluate additional promising candidates such as Nebivolol (delayed ALS progression in SOD1 mice if combined with donepezil)[54], Carvedilol, 20(R)-Propanaxadiol, Lonafarnib (small molecules with previously shown protein aggregation inhibitory effects)[55–57], and Chikusetsusaponin IVa (an autophagy and mitophagy enhancer)[58]. Interestingly, the model appears to prioritise compounds with demonstrated activity against other amyloidogenic proteins (e.g., amyloid-*β*, tau, *α*-synuclein), suggesting that it captures shared chemical scaffolds underlying protein aggregation as well as compounds with broader neuroprotective effects *via* anti-inflammatory, antioxidant, or proteostasis-modulating pathways, reflecting the mechanistic diversity encoded by the model.

Among the top-ranked candidates, berberrubine and PE859 were selected for experimental validation. Molecular docking predicted favourable but distinct interactions with the TDP-43 RNA recognition motif, implying potentially different mechanisms of action. Future work will involve molecular dynamics simulations to evaluate the effect of berberrubine and PE859 binding on RRM stability. Importantly, both compounds significantly reduced TDP-43 aggregation inHEK cells as measured by FLIM. *In vivo* studies in *C. elegans* revealed a significant rescue of TDP-43–induced locomotor defects by PE859, with a partial decrease in locomotion defects for berberrubine.

A key limitation is the heterogeneity of the training data, which combines compounds tested across diverse biological systems, thereby introducing inherent noise. However, as additional data becomes available, both the dataset and model can be further refined. Future work could explore an end-to-end deep learning approach where chemical features and molecular targets are embedded within the graph neural network, with an MLP as the final classifier; here, we selected XGBoost to facilitate interpretability. Overall, this work demonstrates a hybrid machine learning approach for identifying small molecule aggregation inhibitors and could accelerate therapeutic development for TDP-43 proteinopathies and related neurodegenerative diseases.

## 4 Methods

### 4.1 Dataset curation

A dataset of 2,776 small molecules was compiled from ChEMBL (release 35)[22], PubChem[24], and patent records, comprising 294 molecules reported to reduce or protect against TDP-43 aggregation in cells or model organisms (labelled “active”) and 2,482 compounds with no effect or exacerbated aggregation (labelled “inactive”).

### 4.2 Feature generation and pre-processing

For the small molecules in the curated dataset, canonical SMILES strings were obtained from PubChem and further standardised using RDKit’s[21] MolStandardize module (see “Software and implementation” for library versions), including hydrogen removal, metal disconnection, functional group normalisation, fragment selection, charge neutralisation, and tautomer canonicalisation. Chemical descriptors and extended-connectivity fingerprints (ECFPs; number of bits = 2,048, radius = 2)[59] were then generated from the standardised SMILES strings using RDKit. RDKit descriptors were normalised using min-max scaling using scikit-learn[60]. Biological target annotations were extracted from ChEMBL and encoded as binary features indicating whether each compound had been annotated with a given target.

### 4.3 Train–test split and nested cross-validation

The dataset was split into 80% training and 20% test sets using the RDKit Butina ClusterData function to reduce scaffold redundancy[61]. Model performance was assessed using nested 5-fold cross-validation with scikit-learn StratifiedGroupKFold function, where outer folds were grouped by chemical clusters to prevent overestimation from structurally similar compounds.

### 4.4 GNN model training

GNN-derived embeddings were generated using a directed message-passing neural network (D-MPNN) implemented in Chemprop[20]. Standardised SMILES strings were converted to molecular graphs where atoms are nodes and bonds are edges, each represented by feature vectors encoding atomic and bond properties. The D-MPNN architecture represents each covalent bond (e.g., C–O) as two directed edges (e.g., C*→* O and O*→*C), with edge representations iteratively updated by aggregating information from neighbouring bonds over multiple message-passing steps. After message passing, edge embeddings connected to each atom are aggregated into atomic embeddings, which are subsequently aggregated across all atoms to produce a single molecular embedding vector representing the entire compound[20]. These molecular embeddings are then passed through a feedforward neural network for binary classification of molecular activity.

For each outer fold, a separate GNN was trained using scaffold-balanced splitting to further split the outer training fold data into 80% train, 10% validation, 10% test. Hyperparameters were optimised using Ray Tune[62] with HyperOpt search (15 trials). Search ranges for message-passing depth (2–6), feedforward hidden dimensions (300–2400), feedforward layers (1–3), and message hidden dimensions (300–2400) were adapted from Chemprop default example notebooks[20], with dropout (0.1–0.6) and batch size (32 or 64) further added. Models were trained using the Adam optimiser (learning rate = 1*×*10^*−*4^) with binary cross-entropy loss for up to 30 epochs with early stopping (patience=5) based on validation loss, using class weights to address dataset imbalance.

For ensemble models, trained GNNs from each outer fold were used solely for embedding extraction, not prediction, ensuring outer validation sets were never used during GNN training. GNN embeddings were concatenated with RDKit descriptors and ChEMBL annotations to create the final feature set for downstream XGBoost modelling.

### 4.5 Feature selection

Feature selection was performed using scikit-learn within each outer fold to prevent data leakage. ChEMBL target annotations were filtered to retain only targets annotated for at least two compounds in that training fold. Highly correlated features were then removed using a Pearson correlation with a threshold of 80%. Finally, Recursive Feature Elimination (RFE)[27] with an XGBoost estimator[17] was applied to select the most informative features[63]. The numbers of the features selected were 400 for three feature types, 300 for two types, and 200 for single-feature models.

### 4.6 XGBoost model training and hyperparameter optimisation

Following feature selection, XGBoost hyperparameters were optimised within each outer fold using scikit-learn’s 3-fold inner StratifiedGroupKFold cross-validation and RandomizedSearchCV (15 iterations). The XGBoost classifier was configured with eval metric = ‘auc’ and objective = ‘binary:logistic’. The hyperparameter search space included the number of trees (100–400), learning rate (log-uniform, 0.01–0.2), maximum tree depth (3–10), and column subsampling rate (0.1–0.9). Models were optimised using average precision as the primary scoring metric. Performance was evaluated on the outer validation folds using multiple metrics, including Area Under the Receiver Operating Characteristic Curve (ROC-AUC)[64], Matthews correlation coefficient (MCC), F1 score, precision[65], and balanced accuracy[66], implemented with scikit-learn.

### 4.7 Final model training

Following cross-validation, final models were trained on the complete training set. GNN training, embedding extraction, target filtering, correlation filtering, and RFE were repeated on the full training data following the same procedure. Both GNN and XGBoost models were trained using median hyperparameter values from the outer folds and evaluated on the held-out 20% test set.

The final model selected for external validation combined GNN embeddings with RDKit descriptors and ChEMBL target annotations. The optimised GNN parameters were message-passing depth = 3, message hidden dimension = 1300, feedforward hidden dimension = 1500, 3 feedforward layers, and dropout = 0.3, yielding a model with 7.8M total parameters. The XGBoost classifier was configured with n estimators = 300, learning rate = 0.016, max depth = 6, and colsample bytree = 0.4.

### 4.8 Alternative model architectures

Two alternative models were trained for comparison: (i) a standalone GNN using the D-MPNN architecture with embeddings passed directly to a feedforward classifier without extraction or combination with other features, and (ii) a Random Forest[30] classifier (implemented using scikit-learn) combining GNN embeddings, RDKit descriptors, and biological targets. Both followed the same nested cross-validation protocol described above. Random Forest hyperparameter optimisation included n estimators (100–500), max depth (None, 5, 10, 20), max features (‘sqrt’, ‘log2’), and class weight (‘balanced’, None). Additionally, a control model with randomly shuffled training labels was trained using to ensure that performance was not due to chance or data leakage.

### 4.9 Virtual screening of the external dataset

The final, trained model was applied to screen an external library of small molecules compiled from Smer-Barreto *et al*. (2023)[31], comprising of FDA-approved drugs, clinical-stage compounds, and other bioactive molecules from the L2100 TargetMol Anticancer (TargetMol Chemicals, Wellesley Hills, MA) and the L3800 Selleck FDA-approved & Passed Phase (Selleck Chemicals, Houston, TX) libraries. The dataset was pre-processed as described in “Feature generation and pre-processing” and filtered to remove duplicates within the training set as well as internal duplicates with identical standardised SMILES strings, leaving 3,853 small molecules.

TDP-43 anti-aggregation activity probabilities were predicted for all external set molecules. Molecules with predicted probability *>* 80% were subject to further filtering using SwissADME [67] to assess pharmacokinetics and drug-likeness properties. Filtering criteria included: (i) zero violations of Lipinski’s Rule of Five[68], (ii) predicted or literature-reported blood–brain barrier permeability, (iii) zero PAINS[69] and *≤* 1 Brenk[70] structural alerts, and (iv) no prior experimental reports of TDP-43 anti-aggregation activity. From the filtered set of 57 compounds, two were selected for experimental validation, i.e., berberrubine (predicted probability 94%) and PE859 (predicted probability 83%). Tanimoto similarity coefficients (2,048-bit ECFP fingerprints) were calculated using RDKit to evaluate structural similarity to known actives.

### 4.10 Chemical space visualisation

Chemical space was visualised using t-distributed stochastic neighbour embedding (t-SNE) from scikit-learn’s manifold module applied to 2,048-bit ECFP fingerprints. Based on previous work[31], t-SNE was performed with perplexity = 50, max iter = 1,200, learning rate = 200, and init = ‘pca’. Visualisations were displayed using the Python libaries matplotlib and seaborn, and included training set compounds (active and inactive) and external screening hits with predicted probability *>* 80%.

### 4.11 Model interpretability and feature importance analysis

SHAP (SHapley Additive exPlanations)[33] analysis was performed to quantify feature contributions to model predictions, focussing on the top 30 ranked features, of which 15 were GNN-derived embeddings. Pearson correlation coefficients were calculated between top-ranked biological target annotations and chemical descriptors using the pandas Python library to assess feature redundancy and complementarity.

To identify structurally interpretable patterns from the pretrained GNN model, Monte Carlo Tree Search[43] was applied to extract graph-based “rationales”, i.e., molecular substructures associated with increased predicted activity, using Chemprop[20, 59]. Substructures were visualised using RDKit.

### 4.12 Software and implementation

All analyses were performed in Python (v.3.11.13)[71]. Cross-validation and feature selection were implemented using scikit-learn (v.1.7.0)[60], classification using XGBoost (v.2.1.4)[17], while RDKit (v.2025.03.3)[21] was applied for chemical feature generation, Tanimoto similarity calculation, and Butina clustering. Visualisations were created with matplotlib (v.3.10.3)[72] and seaborn (v.0.13.2)[73]. Data manipulation was performed using pandas (v.2.3.0)[74] and numpy (v.2.2.6)[75].

GNN modelling was performed on a high-performance computing cluster using Chemprop (v.2.2.1)[20] with PyTorch (v.2.8.0)[76], PyTorch Lightning (v.2.5.3)[77], and Ray Tune (v.2.49.0) [62]. Models were trained on NVIDIA A100-SXM4-80GB GPU with CUDA 12.8. SHAP analysis was performed using the SHAP library (v.0.48.0)[33]. Reproducibility was ensured by fixing a random seed for all stochastic processes.

### 4.13 Molecular docking

Molecular docking was performed using AutoDock Vina (v.1.2.7)[78]. Protein structures for the N-terminal (X-ray, PDB: 5MDI), NLS (NMR, PDB: 5MRG), and RRM (NMR, PDB: 4BS2) domains were prepared using AutoDock Tools (v.1.5.7)[79]. For NMR structures, the first of the deposited models was used. Protein preparation included solvent removal, polar hydrogen addition, merging of non-polar hydrogens, and Kollman charge[80] computation.

Ligand structures for berberrubine (CID: 72704) and PE859 (CID: 66571561) were obtained from PubChem and prepared with Gasteiger charge[81] assignment in AutoDock Tools. Binding pocket residues were selected based on the work of Rao *et al*. (2021)[46]. Docking was performed with grid spacing of 0.375 Å, generating 9 binding modes with an energy range of 3 kcal/mol. Grid box dimensions were tailored to each binding pocket (28–46 Å per dimension), with exhaustiveness set to 16 for comprehensive conformational sampling. The top-ranked pose by binding affinity was selected for analysis.

Ligand–protein interactions were identified using LigPlot+ (v.2.3)[44], with visualisations created in PyMOL (v.3.1.6)[45]. Manual inspection was performed to identify additional interactions between RRM salt bridge residue Arg151 and PE859.

### 4.14 Compound purchase

Berberrubine chloride (Catalog Number: 15401-69-1) and PE859 (Catalog Number: 1402727-29-0) were obtained from MedChem Express *via* Cambridge Bioscience. Both compounds were dissolved in dimethyl sulphoxide (DMSO; Sigma, Catalog Number: 67-68-5) to reach desired concentrations. Vehicle control wells were treated with 1 *µ*L DMSO for FLIM imaging experiments.

### 4.15 HEK cell culture

HEK-293 cells stably expressing EGFP-TDP-43 (gifted by Professor Christopher Shaw, Institute of Psychiatry, Psychology and Neuroscience, King’s College London) were cultured in Dulbecco’s Modified Eagle Medium:Nutrient Mixture F-12 (DMEM/F-12; ThermoFisher, Catalog number: 11330032), 2 *µ*g/mL Blasticidin (InvivoGen, Catalog number: ant-bl-05), and 10% (v/v) tetracycline-free fetal bovine serum at 37 °C with 5% CO_2_. Cells were maintained in T-25 flasks and passaged twice weekly upon reaching 80% confluency.

### 4.16 Cell viability assay

HEK-293 cells (15,000 per well) were seeded in 96-well plates (Corning, Catalog number: 3603) in 100 *µ*L fresh media. After 24 hours, cells were treated with berberrubine (5, 10 or 50 *µ*M), PE859 (1, 3, 5, 10, or 50 *µ*M), or left untreated (control). Following 24 hours of treatment, 20 *µ*L MTS (CellTiter 96^®^ AQueous One Solution Reagent; Promega, Catalog number: G3580) solution was added to each wells (untreated control, drug-treated and blank wells containing no cells). Cells were incubated at 37 °C with 5% CO_2_ for 3 hours. Absorbance was measured at 490 nm and 650 nm using a microplate reader (CLARIOstar Plus, BMG LABTECH GmbH). Three technical replicates per condition were acquired and averaged. Background-corrected absorbance values were calculated for each well as *Abs*_*corrected*_ = *Abs*_490_ *− Abs*_650_. Cell viability (%) was calculated using the background-corrected absorbance values as shown in using Eq. (1):

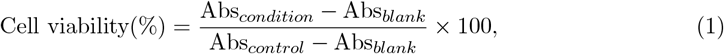

where, *Abs*_*blank*_ represents the mean of blank wells and *Abs*_*control*_ represents the mean of untreated control cells. Data were acquired from four independent biological replicates and analysed in Python (v.3.11.13) using the SciPy (v.1.16.0)[82] and scikit-posthocs (v.0.11.4)[83] libraries. Compounds were selected at concentrations that balanced efficacy with cell viability; specifically, 10 *µ*M berberrubine and 5 *µ*M PE859 were chosen for subsequent experiments.

### 4.17 Fluorescence lifetime imaging microscopy

HEK-293 cells (13,000 per well) were seeded in 8-well Ibidi glass-bottomed plates (Thistle Scientific, Catalog number: 80807) coated with poly-L-lysine (Sigma-Aldrich, Catalog number: P4707) and left to adhere for two days. Cells were then treated with 10 ng/mL doxycycline (Sigma-Aldrich, Catalog number: D9891) for 48 hours to induce EGFP-TDP-43 expression. Following induction, cells were treated with 10 *µ*M berberrubine, 5 *µ*M PE859, or DMSO vehicle control for 24 hours. Cells were maintained at 37 °C with 5% CO_2_ in an incubator system (OKOLab, Italy) during TCSCPC-FLIM imaging.

Imaging was performed on a home-built confocal microscope equipped with a time-correlated single-photon counting (TCSPC) FLIM module, based on an Olympus IX83 inverted microscope with a photomultiplier tube (PMT; PMC-150, Becker & Hickl GmbH, Germany) and an SPC-830 control module (Becker & Hickl GmbH, Germany). Samples were excited with a 40 MHz pulsed supercontinuum laser (Fianium Whitelase, Denmark). EGFP excitation and emission were detected using bandpass filters centred at 474 nm and 542 nm, respectively (FF01–474/27–25 and FF01–542/27–25, Semrock Inc., USA).

Fluorescence lifetime analysis was performed using the phasor plot approach with the FLIMPA software[84]. Data analysis involved cell segmentation by creating manual masks and setting a photon threshold of 100 per pixel in FLIMPA for background removal. Reference fluorescence lifetime correction was conducted by imaging Rhodamine 6G (250 *µ*M in H_2_O) at the beginning of each biological replicate using bandpass filters centered at 510 nm (excitation) and 542 nm (emission) (FF03–510/20–25 and FF01–542/27–25; Semrock Inc., USA). Rhodamine 6G was assigned a reference lifetime of 4 ns for the phasor analysis. Statistical analysis, two-tailed Student’s t-test for berberrubine and Welch’s t-test for PE859, was performed in Python (v.3.11.13) using SciPy (v.1.16.0)[82]. Data represent *≥* 3 independent biological replicates with more than 88 cells analysed per condition.

### 4.18 *C. elegans* strains and culture

The *C. elegans* strains OW1601 (pan-neuronal human TDP-43 expression) and OW1603 (control)[85] were obtained from the Caenorhabditis Genetics Center (CGC). Strains were cultured on nematode growth medium (NGM) prepared as described previously[86]. For drug treatment experiments, NGM plates were supplemented with either 10 *µ*M berberrubine, 10 *µ*M PE859, or no drug addition (control) prior to seeding with live OP50 *E. coli*. Worms were maintained at 16 °C and synchronised through a 6 h egg-laying on plates with corresponding conditions. Synchronised worms were transferred to fresh plates on day 1 of adulthood (young adults), day 3 of adulthood, and day 5 of adulthood, with the locomotion assay performed on day 7 of adulthood.

### 4.19 Swimming assay and behaviour analysis

Worms were transferred to custom-made swimming wells (8mm in diameter) containing 20 *µ*L M9 buffer and recorded using a custom widefield microscope based on an Olympus IX83 frame, equipped with a metal-oxide semiconductor camera (Zyla 4.2, Andor) and controlled by Micro-Manager (Open Imaging)[87]. Recordings were acquired at 30 frames per second using a 1.25× air Olympus objective with dark-field illumination (150 diascopic intensity, 30 ms exposure, 2×2 binning).

The recordings were first pre-processed in Python (v.3.11.13)[71] for background removal and subsequently analysed with CeleST[88] software to extract locomotion parameters. Manual inspection excluded non-moving worms and those not properly tracked by the software. Data were acquired over *≥*3 biological replicates, with 33 control (OW1603), 34 TDP-43 (OW1601), 34 TDP-43 + berberrubine (OW1601), and 34 TDP-43 + PE859 (OW1601) worms analysed.Statistical analysis were performed using the Mann–Whitney U test for body wave number and Student’s t-test for travel speed with SciPy (v.1.16.0)[82].

## 5 Supplementary Information

- Supplementary File 1 (.docx): Supplementary Tables 1-2 and Supplementary Figures 1-3.

## 6 Competing interests

Authors declare no competing interests.

## Acknowledgements

G.S.K.S. acknowledges funding from the Wellcome Trust (065807/Z/01/Z) (203249/Z/16/Z), the UK Medical Research Council (MRC) (MR/K02292X/1), Alzheimer Research UK (ARUK) (ARUK-PG013-14), Michael J Fox Foundation (16238 and 022159), and Infinitus China Ltd.C.F.K. acknowledges funding from the UK Engineering and Physical Sciences Research Council (EP/L015889/1 and EP/H018301/1), the Wellcome Trust (3-3249/Z/16/Z and 089703/Z/09/Z), the UK Medical Research Council (MR/K015850/1 and MR/K02292X/1), and Infinitus China Ltd. S.K. acknowledges funding from AstraZeneca and the UK Engineering and Physical Sciences Research Council (EPSRC) grant EP/S023046/1 for the Centre for Doctoral Training in Sensor Technologies for a Healthy and Sustainable Future. N.F.L. acknowledges the Swiss National Science Foundation (Grant Number P2EZP2 199843). S.V. acknowledges UK Research and Innovation and University of Cambridge. *C. elegans* strains were provided by the CGC, which is funded by NIH Office of Research Infrastructure Programs (P40 OD010440).

## Notes

### Competing Interest Statement

The authors have declared no competing interest.

