## Supplementary Tables 1-2 and Supplementary Figures 1-3 for "Discovery of TDP-43 aggregation inhibitors *via* a hybrid machine learning framework"

**Supplementary Table 1. Performance metrics on 5-fold nested cross-validation for models trained using individual and combined feature types**. Random shuffling of target values (Y shuffle) served as a control to verify that performance was not attributable to information leakage.

| Features | Classifier | ROC-AUC | MCC | F1-score | Balanced Accuracy | Precision |
|---|---|---|---|---|---|---|
| GNN embeddings + chemical descriptors + biological targets | XGBoost | 0.84 ± 0.03 | **0.33 ± 0.04** | **0.30 ± 0.06** | **0.59 ± 0.02** | 0.73 ± 0.06 |
| ECFPs (2048) chemical descriptors + biological targets | XGBoost | **0.87 ± 0.02** | 0.30 ± 0.07 | 0.22 ± 0.08 | 0.56 ± 0.03 | **0.84 ± 0.12** |
| chemical descriptors + biological targets | XGBoost | **0.87 ± 0.01** | 0.28 ± 0.07 | 0.22 ± 0.11 | 0.57 ± 0.03 | 0.78 ± 0.16 |
| biological targets | XGBoost | 0.85 ± 0.03 | 0.21 ± 0.12 | 0.18 ± 0.13 | 0.55 ± 0.04 | 0.55 ± 0.17 |
| chemical descriptors | XGBoost | 0.74 ± 0.05 | 0.07 ± 0.10 | 0.03 ± 0.03 | 0.51 ± 0.01 | 0.50 ± 0.50 |
| GNN network | MLP | 0.70 ± 0.05 | 0.06 ± 0.08 | 0.03 ± 0.04 | 0.51 ± 0.01 | 0.30 ± 0.45 |
| ECFPs (2048) | XGBoost | 0.68 ± 0.04 | 0.10 ± 0.1 | 0.12 ± 0.08 | 0.53 ± 0.03 | 0.28 ± 0.17 |
| random Y shuffle (GNN embeddings + chemical descriptors + biological targets) | XGBoost | 0.50 ± 0.05 | -0.01 ± 0.01 | 0.00 ± 0.00 | 0.50 ± 0.00 | 0.00 ± 0.00 |
| GNN embeddings + chemical descriptors + biological targets | Random Forest | 0.81 ± 0.04 | 0.17 ± 0.09 | 0.14 ± 0.07 | 0.54 ± 0.02 | 0.54 ± 0.23 |

**Supplementary Table 2. Monte Carlo Tree Search-identified graph-based rationales associated with increased predicted for test set compounds correctly predicted as active.** Rationales are highlighted in blue.

| | | |
|---|---|---|
| MLS003120626 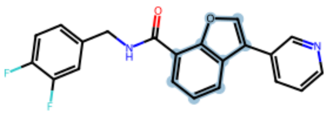 | 4-Chloro-7-methoxy-5H-pyrimido[5,4-b]indole 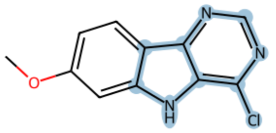 | (2z)-N-(2,5-Dichlorophenyl)-2-(Hydroxyimino)-3,4-Dihydro-2h-1-Benzopyran-3-Carboxamide 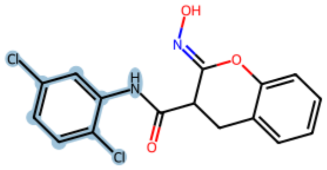 |
| SMR001277452 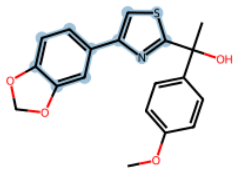 | MLS003120491 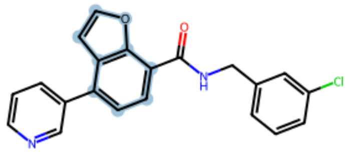 | N-[2-(4-methoxyphenyl)quinolin-4-yl]-N',N'-dimethylpropane-1,3-diamine 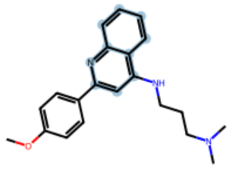 |
| MLS003120620 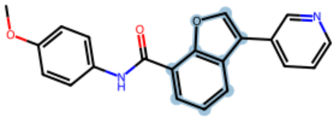 | 3-bromo-N-(2,3-dihydro-1,4-benzodioxin-6-ylcarbamothioyl)benzamide 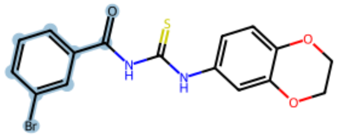 | N-(1,3-benzothiazol-2-yl)-2-thiophen-2-ylacetamide 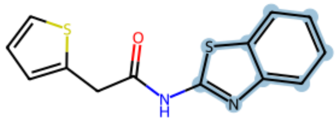 |
| MLS001060488 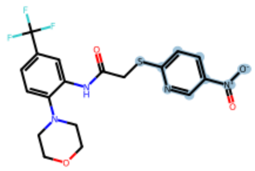 | 338748-09-7 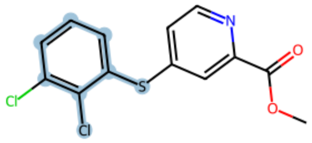 | SMR000005287 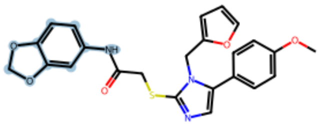 |

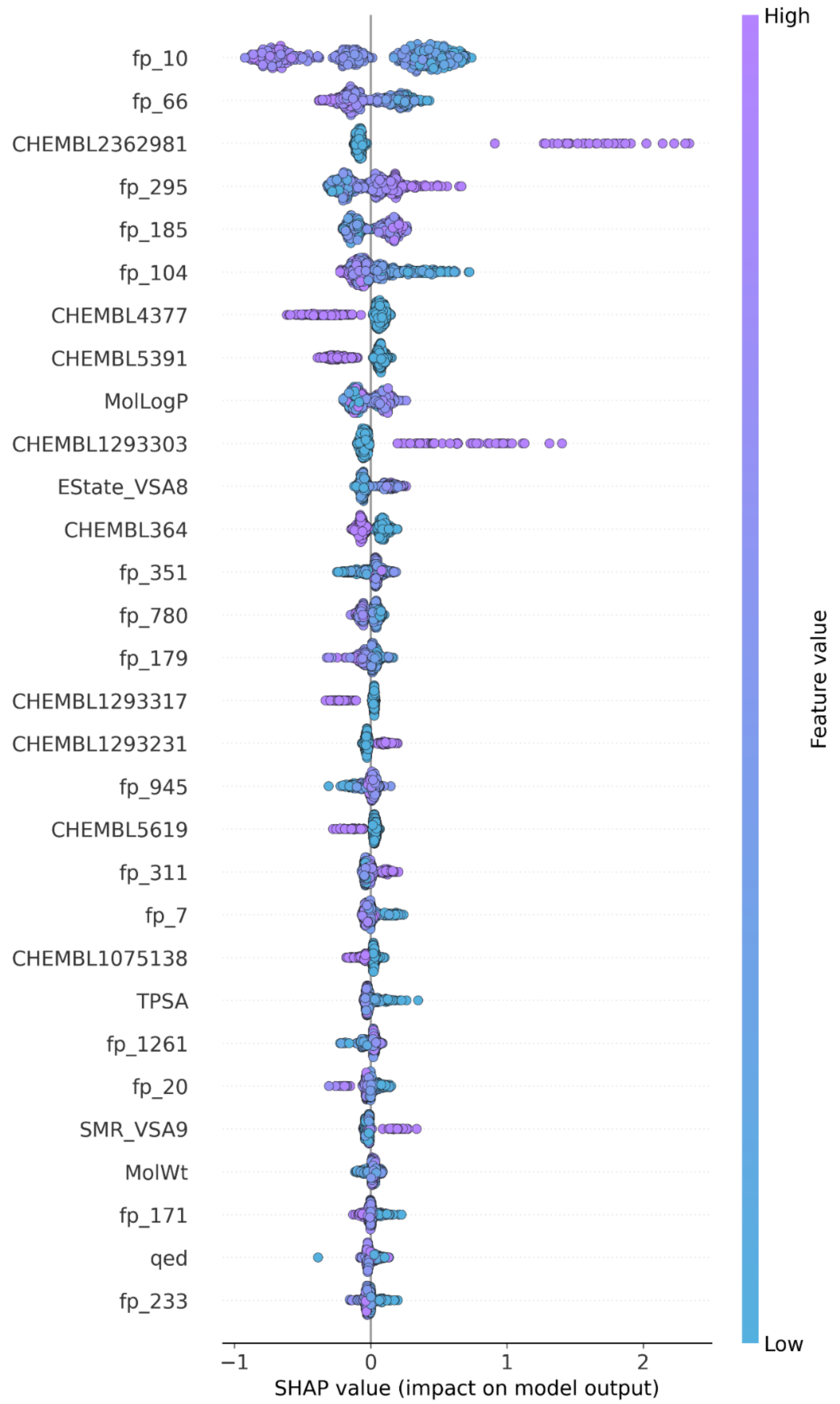

**Supplementary Figure 1. Top performing features (n= 30) of the XGBoost model trained using GNN embeddings, RDKit chemical descriptions, and ChEMBL targets, identified using SHAP analysis**. Half of the top-ranked descriptors correspond to GNN embeddings denoted by fp_. Points show individual compounds coloured by the corresponding SHAP value for that feature, where purple indicates a high SHAP value and blue represents a low one.

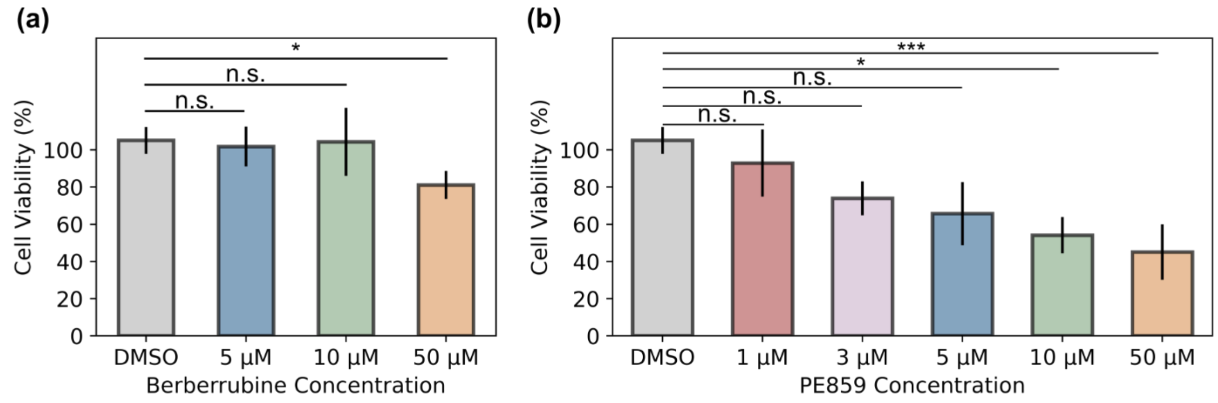

**Supplementary Figure 2. Cell viability assessed by MTS assay following treatment with increasing concentrations of (a) berberrubine and (b) PE859 in HEK cells**. Data were acquired from four biological replicates. Statistical comparisons were performed using one-way ANOVA with Tukey's HSD (berberrubine; normal distribution) or Kruskal–Wallis with Bonferroni-corrected Dunn's test (PE859; non-normal distribution). *$p < 0.05$, **$p < 0.01$, ***$p < 0.001$; n.s., non-significant.

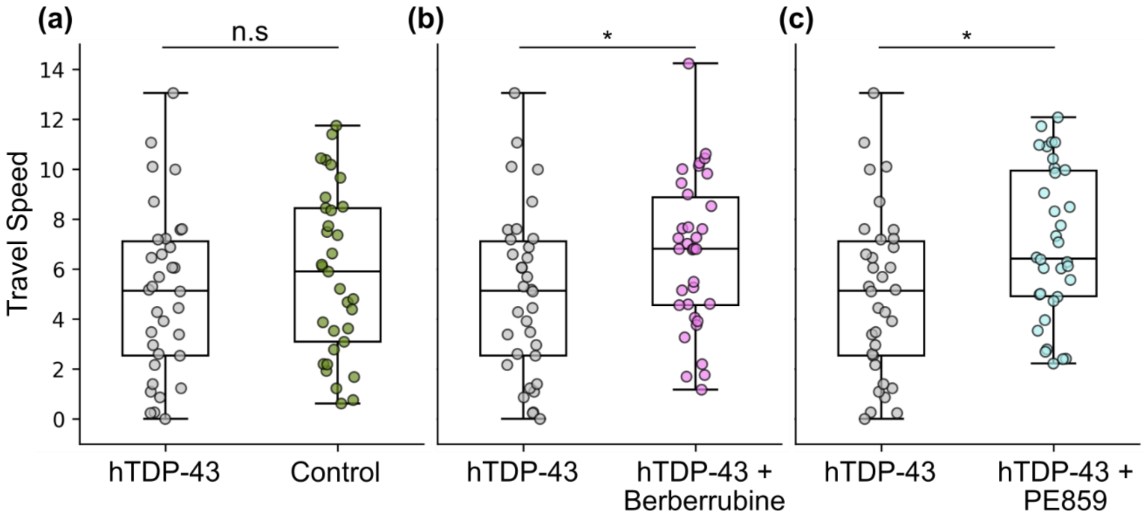

**Supplementary Figure 3. Treatment with candidate compounds significantly increases travel speed of hTDP-43 worms.** (a) Control worms display modestly higher locomotion speed compared to hTDP-43 worms, though this difference is not significant. Treatment with (b) berberrubine and (c) PE859 both significantly increase travel speed in worms expressing hTDP-43 pan-neuronally compared to untreated hTDP-43 worms. Statistical analysis was performed using Student's t-test, with *$p < 0.05$ and n.s. denoting not significant.
